# Cube-based screening identifies a quinoa-derived synthetic microbial community that promotes plant growth and modulates root epidermal responses under salt stress

**DOI:** 10.64898/2026.07.15.738596

**Authors:** Hatairat Dangjarean, Yoshinori Murata, Yasufumi Kobayashi, Sara Neyrot, Takuya Ogata, Yasunari Fujita

**Author notes:** Corresponding author: Yoshinori Murata, Yasunari Fujita.

## Abstract

Plant-associated bacteria can improve plant performance under abiotic stress, but beneficial functions in plant microbiomes may depend on defined combinations of microorganisms rather than individual isolates alone. Here, we developed a cube-based screening strategy to identify functional synthetic microbial communities (SynComs) from 135 quinoa-associated bacterial isolates while preserving combinatorial diversity and traceability of isolate-level contributions. The isolates were divided into five 27-isolate sets, each arranged as a 3 × 3 × 3 cube in which each 3 × 3 layer was defined as a 9-isolate SynCom, generating 45 SynComs in total. Screening under 100 mM NaCl identified SynCom DY1 (SCDY1) as a candidate salt stress-mitigating consortium. SCDY1 consisted of nine taxonomically diverse isolates and exhibited a multifunctional profile, including siderophore production, phosphate solubilization, carboxymethyl cellulose degradation, indole compound production, and growth under saline conditions. In *Arabidopsis thaliana*, SCDY1 promoted primary root elongation and biomass accumulation in a salinity-dependent manner, with the clearest effect under 120 mM NaCl, and at least a subset of constituent bacteria was recoverable from inoculated seedlings. RNA sequencing and targeted RT-qPCR indicated that SCDY1 modulated host gene expression under moderate salinity stress, with responsive genes associated with oxidative stress, water- and oxygen-related processes, phenylpropanoid biosynthesis, glutathione metabolism, and root epidermis-related processes. Root hair phenotyping further showed that SCDY1 enhanced root hair-related traits and shifted visible root hair formation closer to the root apex. These findings identify a quinoa-derived SynCom that improves plant performance under salinity stress and provide a practical, traceable framework for discovering beneficial microbial consortia from plant-associated bacterial collections.

**Scope statement:** This manuscript fits the Research Topic “Harnessing Plant Microbiomes for Climate Resilience: From Ecological Insight to Synthetic Community Design” in Frontiers in Plant Science because it presents a traceable strategy for discovering functional synthetic microbial communities from a stress-adapted plant-associated bacterial collection. We developed a cube-based screening strategy using 135 quinoa-associated bacterial isolates and identified a nine-isolate synthetic microbial community, SCDY1, that promotes *Arabidopsis* growth under moderate salinity stress. The study integrates microbiological screening, characterization of plant growth-promoting traits, bacterial re-isolation, plant growth phenotyping, RNA-seq, RT-qPCR, and root hair phenotyping. These analyses link SCDY1 treatment to salinity-dependent growth promotion, recoverable bacterial members, stress- and redox-associated transcriptional changes, phenylpropanoid-related responses, and modulation of root epidermal phenotypes. By connecting a defined SynCom with host transcriptional and root epidermal responses, this work advances understanding of beneficial plant–microbe interactions under salt stress. The cube-based design also provides a practical and traceable framework for discovering functional SynComs from large plant-associated bacterial collections, which should be of interest to researchers studying plant symbiosis, microbiome engineering, abiotic stress tolerance, and sustainable crop improvement.

## 1 Introduction

Soil salinity is a major environmental constraint limiting plant growth and agricultural productivity worldwide (Munns and Tester, 2008; Tang et al., 2024). Under saline conditions, plants are exposed to osmotic stress, ionic imbalance, nutrient disturbance, and oxidative stress, which collectively restrict root development, biomass accumulation, and yield (Hossain and Dietz, 2016; Munns and Tester, 2008). Because salinization is accelerated by irrigation, soil degradation, and climate change, sustainable strategies for improving plant performance under saline environments are increasingly needed (Tang et al., 2024). In addition to plant breeding and agronomic management, plant-associated microorganisms have attracted attention as biological resources that may help plants cope with abiotic stress.

Plant growth-promoting bacteria can influence host performance through diverse mechanisms, including phytohormone production, nutrient mobilization, siderophore production, modulation of stress responses, and alteration of root system architecture (Backer et al., 2018; Lugtenberg and Kamilova, 2009; Vacheron et al., 2013). Several bacterial isolates have been reported to promote plant growth under salt stress, indicating that plant-associated bacteria can contribute to stress adaptation through bacterial physiological functions as well as host transcriptional and physiological responses (Etesami and Glick, 2024; Yang et al., 2009). However, in natural plant-associated environments, bacteria rarely act as isolated entities. Instead, plant microbiomes consist of multiple interacting taxa whose collective functions may differ from those predicted from single-isolate traits alone (Trivedi et al., 2020). This complexity may partly explain why single-isolate inoculants can show context-dependent performance across plant species, stress intensities, and growth environments (Backer et al., 2018; Trivedi et al., 2020).

Synthetic microbial communities (SynComs) provide a powerful approach for studying and utilizing microbiome functions under controlled and reproducible conditions (Johns et al., 2016; Trivedi et al., 2020; Vorholt et al., 2017). Because SynComs are composed of defined microbial members, they offer greater reproducibility and interpretability than undefined microbial communities while retaining selected community-level properties. SynComs have been constructed using several strategies, including simplification of natural microbiomes, selection of core community members, trait-based selection of functionally characterized isolates, and bottom-up assembly of defined microbial consortia. These approaches have advanced both plant microbiome research and broader synthetic community studies, but the design of effective SynComs remains challenging because community functions cannot always be predicted from individual isolate traits alone. In principle, combinatorial screening can reveal unexpected beneficial combinations; however, exhaustive testing rapidly becomes impractical as isolate number increases. Conversely, purely trait-based or fully rational designs may overlook beneficial interactions that emerge only in specific combinations. Thus, a practical challenge in SynCom-based approaches is how to preserve sufficient combinatorial diversity to discover unexpected community functions while retaining enough structure to infer whether beneficial activity is associated with dominant individual isolates or with specific combinations of multiple isolates. Previous studies using microbiomes derived from stress-adapted plants, including halophytes and desert plants, have demonstrated that selected microbial communities can improve the performance of non-host plants under salinity stress (Schmitz et al., 2022; Yuan et al., 2016). These studies support the idea that stress-adapted plants are valuable sources of beneficial microorganisms and highlight the need for screening strategies that can efficiently identify functional SynComs from large isolate collections. Plant-associated bacterial communities commonly include members of major phyla such as Pseudomonadota/Proteobacteria, Bacillota/Firmicutes, Actinomycetota/Actinobacteria, and Bacteroidota/Bacteroidetes (Choi et al., 2021). Therefore, SynCom screening strategies based on large isolate collections should ideally preserve broad taxonomic representation while remaining experimentally tractable.

Quinoa (*Chenopodium quinoa* Willd.) is a stress-resilient crop originating from the Andean region and is recognized for its broad adaptation to harsh environments, including saline and drought-prone conditions (Bazile et al., 2016; Hariadi et al., 2011). Recent genomic and transcriptomic studies have advanced our understanding of quinoa stress adaptation and salt tolerance, including genotype-dependent variation in salt-responsive growth traits, Na□ exclusion mechanisms, and the expansion of chromosome-scale genome resources across diverse quinoa accessions (Jarvis et al., 2017; Kobayashi et al., 2024; Kobayashi et al., 2025; Mizuno et al., 2020; Rey et al., 2024; Rey et al., 2023; Yasui et al., 2016). In parallel, virus-induced gene silencing and virus-mediated overexpression systems have enabled functional analyses of quinoa genes, further strengthening quinoa as an emerging model for stress-resilient crop research (Ogata et al., 2021). Given its adaptation to stressful environments and the availability of genomic and functional resources, quinoa may represent a promising source of stress-associated microorganisms with potential plant growth-promoting functions. We previously identified a quinoa-associated bacterial isolate that promoted plant growth specifically under salinity stress conditions and exhibited salinity-responsive bacterial physiological traits (Murata et al., 2026). This single-isolate analysis suggested that quinoa-associated bacteria can contribute to plant–microbe interactions under salt stress. However, it remained unclear whether beneficial functions in quinoa-associated bacterial collections are primarily attributable to individual isolates or to combinations of multiple isolates.

In the present study, we developed a cube-based SynCom screening strategy to identify salt stress-mitigating bacterial consortia from quinoa-associated isolates. This strategy was designed to preserve structured combinatorial diversity while keeping the number of SynComs within a practical screening scale. By arranging isolates in defined cubes, each isolate was included in multiple SynComs, allowing potential isolate-level contributions to remain traceable while testing different isolate combinations. Using this approach, we identified SynCom DY1 (SCDY1) as a quinoa-derived consortium that promotes *Arabidopsis* growth under moderate salinity stress. We further characterized its plant growth-promoting traits, recoverability from inoculated seedlings, and associated host transcriptional and root epidermal responses. Together, this study provides a practical framework for discovering beneficial SynComs from plant-associated bacterial collections and highlights quinoa-associated bacterial communities as promising resources for improving plant performance under salinity stress.

## 2 Materials and methods

### 2.1 Isolation and taxonomic identification of quinoa-associated bacterial isolates

Endophytic bacteria were isolated from seeds and seedlings of Chenopodium quinoa Willd. The Kd inbred line (Yasui et al., 2016) was used as the main source material, and all seedling-derived isolates were obtained from Kd seedlings. Commercially available seeds derived from southern highland-type quinoa were also used for seed-borne bacterial isolation. The isolation procedure was performed as described previously (Murata et al., 2026), with minor modifications.

For isolation of seed-borne bacteria, quinoa seeds were surface-sterilized with 70% ethanol for 15–30 s or with a 1:100 dilution of Plant Preservative Mixture (Plant Cell Technology, Inc., Washington, DC, USA) for up to 5 min, followed by four rinses with sterile Milli-Q water. The sterilized seeds were imbibed in sterile Milli-Q water and placed on Reasoner’s 2A (R2A) agar plates or trypticase soy agar (TSA) plates. R2A agar plates were prepared with Difco R2A Agar at 18.2 g/L, and TSA plates were prepared with 3% (w/v) Trypticase Soy Broth and 1.8% (w/v) Difco Agar. All media components were purchased from Becton, Dickinson and Company (Franklin Lakes, NJ, USA). Plates were incubated at 25°C for 2–3 days, and bacterial colonies emerging from the seeds were isolated and purified. To confirm surface sterilization, the final wash solution was spread on R2A agar plates and incubated at 25°C.

For isolation of seedling-derived bacteria, quinoa seeds were surface-sterilized with 70% ethanol for 5–10 min, followed by immersion in 1.0% sodium hypochlorite solution for 5–10 min, and subsequently rinsed four times with sterile Milli-Q water. The sterilized seeds were placed on half-strength Murashige and Skoog (0.5× MS) agar medium supplemented with 3% (w/v) sucrose and incubated at 25°C until germination. The medium was prepared with 0.5× MS salts (Fujifilm Wako Pure Chemical Corporation, Osaka, Japan) and 0.8% (w/v) Difco Agar (Becton, Dickinson and Company). Six days after germination, quinoa seedlings were transferred to cell trays containing a sterilized substrate composed of vermiculite and Potting Mix for Professionals (Innovex, Tokyo, Japan) at a ratio of 9:1 (v/v). The substrate had been autoclaved and dried before use. Seedlings were watered for one day and subsequently cultivated at 25°C for 7 days under salt stress conditions with 300 mM NaCl.

After salt treatment, seedlings were surface-sterilized by washing three times with sterile Milli-Q water, followed by immersion in 70% ethanol for 1 min, immersion in 1.0% sodium hypochlorite solution for 1 min, and two additional washes with sterile Milli-Q water. To confirm the effectiveness of this final surface-sterilization step, the final rinse water was spread onto R2A agar plates and incubated at 25°C for 3 days; no colony formation was observed. Residual moisture was removed using sterile paper towels. The sterilized tissues were homogenized with sterile zirconia beads at 1,100 rpm for 3 min using a Shake Master homogenizer (BMS-A20TP, BioMedical Science, Tokyo, Japan). The resulting homogenates were spread onto TSA and R2A agar plates and incubated at 25°C for 4 days. Distinct bacterial colonies that appeared on the plates were isolated and purified.

Genomic DNA was extracted from each bacterial isolate using the Wizard Genomic DNA Purification Kit (Promega, Madison, WI, USA) according to the manufacturer’s instructions. Partial 16S rRNA gene sequences were amplified by PCR using MightyAmp DNA Polymerase Ver. 3.0 (Takara Bio Inc., Shiga, Japan) with the universal primers 27F and 1492R. PCR products were purified using the Wizard SV Gel and PCR Clean-Up System (Promega). The obtained sequences were compared with reference sequences using nucleotide BLAST searches, and taxonomic assignments were made based on the closest sequence similarities. A total of 135 quinoa-associated bacterial isolates were selected for SynCom construction. Based on partial 16S rRNA gene sequences, the 135 isolates were classified into four bacterial phyla: Bacillota (formerly Firmicutes; 57 isolates), Pseudomonadota (formerly Proteobacteria; 56 isolates), Actinomycetota (formerly Actinobacteria; 16 isolates), and Bacteroidota (formerly Bacteroidetes; 6 isolates). The taxonomic classification and cube-based allocation of the 135 isolates are listed in Supplementary Table 1. Isolate IDs indicate the original identification numbers assigned to the bacterial isolates.

### 2.2 Cube-based construction and screening of synthetic microbial communities

To construct defined synthetic microbial communities, the 135 bacterial isolates were divided into five independent sets of 27 isolates. The sets were assembled to include representatives of major plant-associated bacterial phyla and to maintain broad taxonomic diversity while avoiding overlap among isolates. During SynCom construction, isolates that did not grow reliably or could not be handled consistently under the experimental conditions were excluded or replaced. Although the taxonomic composition differed among the five sets, broad taxonomic representation was retained across the screening collection. Each 27-isolate set was arranged in a 3 × 3 × 3 cube. In each cube, every 3 × 3 layer along the x-, y-, and z-axes was defined as one 9-isolate SynCom, generating nine SynComs per cube and 45 SynComs in total. In this design, each isolate was included in three SynComs, one in each axial direction. The composition of the 45 SynComs generated by this cube-based design is listed in Supplementary Table 2.

For SynCom preparation, each constituent isolate was cultured in trypticase soy broth (TSB; 3% w/v) at 25°C with shaking at 150 rpm, except for isolate 91, which was cultured at 91 rpm because it did not grow sufficiently at 150 rpm. To reduce the influence of differences in growth rate among isolates, bacterial cultures at different growth stages were combined before SynCom assembly. Each culture was adjusted to OD600 = 0.5, and the nine constituent isolates were mixed in equal volumes. The mixed bacterial suspension was washed at least three times with 10 mM MgCl□ to remove residual medium and then resuspended in 10 mM MgCl□. The final SynCom suspension was adjusted to OD600 = 0.025 unless otherwise stated. Mock treatment consisted of 10 mM MgCl□ without bacteria.

The 45 SynComs were screened for their ability to mitigate salt stress-associated growth inhibition in *Arabidopsis thaliana* Col-0 (CS60000). For primary screening, *Arabidopsis* seeds were sown on nylon mesh placed on the surface of rectangular agar plates. Nylon mesh sheets (NB34, nylon 66PA, 512-µm opening; Tokyo Screen Co., Ltd., Tokyo, Japan) were cut to 12 × 8 cm, soaked in 70% ethanol for 30 min, and UV-sterilized in a laminar flow cabinet for 30 min on each side. The sterilized mesh was placed on rectangular plates (10 × 14 cm) containing 65 mL of agar medium composed of 0.5× MS salts (Fujifilm Wako Pure Chemical Corporation, Osaka, Japan), 3% (w/v) sucrose, 0.05% (w/v) MES-KOH (pH 5.7), and 0.8% (w/v) agar. Sixty *Arabidopsis* seeds were sown on each mesh-covered plate. Plates were incubated horizontally at 22°C under long-day conditions (16 h light/8 h dark) without cold stratification, and their positions were rotated daily to minimize positional effects.

At 5–6 days after sowing, seedlings were transferred to screening plates by moving the entire nylon mesh with sterile forceps. Screening plates were rectangular plates (10 × 14 cm) containing 65 mL of agar medium composed of 0.5× MS salts, 0% (w/v) sucrose, 0.05% (w/v) MES-KOH (pH 5.7), and 0.8% (w/v) agar, with or without 100 mM NaCl. Before transfer, 1 mL of SynCom suspension at OD600 = 0.025 or 10 mM MgCl□ mock solution was spread evenly over the surface of each screening plate using a sterile spreader and allowed to dry. For each 9-isolate SynCom, seedlings were transferred to two replicate plates without NaCl and two replicate plates containing 100 mM NaCl. The primary screening experiment was performed twice on independent days.

SynCom performance was evaluated based on salinity-dependent growth promotion. SynComs showing little or no effect under 0 mM NaCl but improved seedling growth under 100 mM NaCl relative to mock-treated controls were selected as primary candidates. Eleven SynComs met this criterion and were further assessed by leaf area measurement using ImageJ/Fiji. Among these candidates, SynCom DY1 showed the most consistent and reproducible positive response under 100 mM NaCl and was selected for subsequent analyses as SCDY1. The closest related taxa and sequence-based taxonomic information for the SCDY1 constituent isolates are listed in Supplementary Table 3.

### 2.3 Phylogenetic analysis of SCDY1 constituent isolates

The nine bacterial isolates constituting SCDY1 were subjected to phylogenetic analysis based on partial 16S rRNA gene sequences. Sequences of SCDY1 constituent isolates and closely related reference taxa were aligned using MUSCLE. A phylogenetic tree was constructed using the neighbor-joining method with 1,000 bootstrap replicates. The resulting tree was used to examine the taxonomic relationships of the nine SCDY1 constituent isolates.

### 2.4 Characterization of plant growth-promoting traits and bacterial growth under saline conditions

Plant growth-promoting traits of SCDY1 and its individual constituent isolates were characterized using plate-based and colorimetric assays. The tested traits included siderophore production, phosphate solubilization, carboxymethyl cellulose degradation, indole compound production, and bacterial growth under saline conditions. For plate-based assays, bacterial suspensions of SCDY1 or each individual constituent isolate were prepared in 10 mM MgCl□ and adjusted to the indicated optical density. Aliquots of 3–4 µL were spotted onto the corresponding assay medium and incubated at room temperature. Plates were scanned using a flatbed scanner, and colony diameters, halo diameters, or activity zones were quantified using ImageJ/Fiji.

Siderophore production was evaluated using chrome azurol S (CAS) agar (Louden et al., 2011; Murata et al., 2026), with minor modifications. Bacterial suspensions were spotted onto CAS agar plates, and siderophore production was assessed based on the formation of orange halo zones around bacterial colonies. Thirteen days after inoculation, colony diameter and the diameter of the surrounding orange halo were measured. Phosphate solubilization activity was evaluated using phosphate agar medium containing tricalcium phosphate as an insoluble phosphate source (Chen and Liu, 2019; Murata et al., 2026), with minor modifications. Bacterial suspensions were spotted onto phosphate agar plates and incubated for 13 days. Phosphate solubilization activity was evaluated by measuring the clear halo diameter surrounding bacterial colonies. Carboxymethyl cellulose (CMC) degradation activity was evaluated using CMC-Na agar medium (Liang et al., 2014; Murata et al., 2026), with minor modifications. Bacterial suspensions were spotted onto CMC-Na agar plates and incubated for 7 days. Plates were subsequently stained with 0.1% (w/v) Congo-red solution, and the clear halo diameter surrounding bacterial colonies was measured as an indicator of CMC degradation activity. Indole compound production was quantified using the Salkowski reagent method (Gordon and Weber, 1951). Bacterial cultures were grown in 1/10-strength TSB supplemented with 1 mM L-tryptophan, and culture supernatants were mixed with Salkowski reagent. After incubation in the dark at room temperature, absorbance was measured at 530 nm. Indole compound concentration was estimated using a standard curve prepared with indole-3-acetic acid. Bacterial growth under saline conditions was evaluated by monitoring optical density at 600 nm over 50 h in culture medium supplemented with 0, 100, 120, or 150 mM NaCl. Multifunctional profiles of SCDY1 and its constituent isolates were visualized using radar plots generated with the fmsb package in R.

### 2.5 *Arabidopsis* growth assay under salinity stress

*Arabidopsis thaliana* Col-0 (CS60000) seeds were surface-sterilized and stratified at 4°C for 48 h to synchronize germination (Nagatoshi et al., 2023). Seeds were germinated on modified 0.5× MS medium containing 3% sucrose and agar. Seedlings were grown vertically in a controlled growth chamber at 22°C under long-day conditions with white light at an intensity of 45 µmol m□² s□¹ and 60% relative humidity. Uniformly sized seedlings were transferred to 0.5× MS medium without sucrose and supplemented with or without NaCl. For the initial growth assay, seedlings were transferred to medium containing 0 or 100 mM NaCl. After acclimation, *Arabidopsis* seedlings were inoculated with 4 µL of SCDY1 suspension adjusted to A600 = 0.025 near the root tip region. A 10 mM MgCl□ solution was applied in the same manner as the mock control. Plates were maintained vertically, and their positions were rotated daily to minimize positional effects. Primary root length was monitored over time by marking root tips and scanning plates with a flatbed scanner. Root length was quantified from scanned images using ImageJ/Fiji. Fresh weight was measured immediately after harvesting seedlings. Dry weight was determined after drying plant samples at 60°C until a constant weight was obtained.

To further evaluate the salinity range of SCDY1 activity, additional assays were performed under 0, 120, and 150 mM NaCl conditions. Seedlings with similar primary root lengths were transferred to the corresponding salinity media and acclimated before inoculation. SCDY1 suspension or 10 mM MgCl□ mock solution was applied to each seedling, and plants were grown vertically under controlled conditions. After treatment, primary root length, fresh weight, and dry weight were measured. The experiment was independently repeated to confirm reproducibility.

### 2.6 Re-isolation of SCDY1-derived bacteria from inoculated seedlings

To examine whether SCDY1-derived bacteria could be recovered from inoculated plants, *Arabidopsis thaliana* seedlings were transferred to 0.5× MS medium supplemented with 100, 120, or 150 mM NaCl (Murata et al., 2026), with minor modifications. One day after transfer, seedlings were inoculated with 4 µL of SCDY1 suspension adjusted to OD_600_ = 0.025 or with 4 µL of 10 mM MgCl□ as the mock control. Six days after inoculation, plant tissues were homogenized with sterile zirconia beads at 1,100 rpm for 3 min using a Shake Master homogenizer, and serial dilutions of the homogenates were spread onto tryptic soy agar, one-tenth-strength tryptic soy agar, and R2A agar plates. Bacterial colonies recovered from SCDY1-inoculated seedlings were subjected to colony PCR and sequence analysis for taxonomic identification. The re-isolation experiment was performed twice independently.

### 2.7 RNA extraction and RNA-seq analysis

For transcriptome analysis, *Arabidopsis* seedlings were grown under four treatment conditions: mock-treated seedlings under non-saline conditions, mock-treated seedlings under 120 mM NaCl, SCDY1-treated seedlings under non-saline conditions, and SCDY1-treated seedlings under 120 mM NaCl. These groups were designated M0, M120, S0, and S120, respectively. SCDY1 or mock solution was applied to seedlings, and plant samples were collected at 100 h after inoculation, when SCDY1-dependent growth differences were detectable under 120 mM NaCl. Four biological replicates were prepared for each treatment, with each replicate consisting of pooled seedlings.

Total RNA was extracted using RNAiso Plus (Takara Bio Inc., Shiga, Japan) according to the manufacturer’s protocol (Nagatoshi et al., 2023), with minor modifications. RNA quality and quantity were assessed before library preparation. RNA-seq libraries were constructed and sequenced using the Illumina NovaSeq platform (Illumina Inc., San Diego, CA, USA). Raw sequencing reads were quality-filtered to remove adapter sequences and low-quality bases using Trimmomatic v0.41 (Bolger et al., 2014). Cleaned reads were aligned to the *Arabidopsis thaliana* TAIR10 reference genome using HISAT2 v2.1.0 (Kim et al., 2019). Gene-level read counts were quantified using featureCounts (Liao et al., 2014), and gene expression levels were calculated as transcripts per million. Differentially expressed genes were identified using edgeR (Robinson et al., 2010). Four pairwise comparisons were analyzed: M120 vs M0, S120 vs S0, S0 vs M0, and S120 vs M120. Genes with an absolute log□ fold change of at least 0.585 and a false discovery rate below 0.05 were defined as differentially expressed genes. DEG distributions were visualized using bar plots and volcano plots. Principal component analysis was performed using log□-transformed expression values. Overlaps among DEG sets were visualized using Venn diagrams.

### 2.8 Functional enrichment analysis

Gene Ontology enrichment analysis was performed for selected DEG sets to identify enriched biological processes and molecular functions. GO enrichment analysis was performed using the clusterProfiler package (Yu et al., 2012) with the Arabidopsis annotation package org.At.tair.db. Enrichment significance was evaluated using Benjamini–Hochberg-adjusted *p* values (Benjamini and Hochberg, 1995). SCDY1-specific DEGs under salinity stress were defined as DEGs uniquely identified in the S120 vs M120 comparison among the four pairwise comparisons. KEGG pathway enrichment analysis was performed using DEGs identified in the S120 vs M120 comparison to identify metabolic and signaling pathways associated with SCDY1-responsive transcriptional changes under salinity stress (Kanehisa and Goto, 2000). Enriched KEGG pathways were identified using adjusted *p* values, and selected enriched pathways were further visualized using gene expression heatmaps.

### 2.9 RT-qPCR analysis

To confirm selected RNA-seq expression patterns using a targeted quantification method, RT-qPCR analysis was performed using the same RNA samples that were used for RNA-seq analysis from mock-treated and SCDY1-treated *Arabidopsis* seedlings grown under 120 mM NaCl. Total RNA was extracted from plant tissues using RNAiso Plus. For first-strand cDNA synthesis, 1 µg of total RNA was treated with RQ1 RNase-Free DNase (Promega, Madison, WI, USA) for 30 min at 37°C, and cDNA was synthesized using the SuperScript III First-Strand Synthesis System (Invitrogen, Carlsbad, CA, USA) according to the manufacturer’s instructions. RT-qPCR was performed on a QuantStudio 7 Flex Real-Time PCR System (Applied Biosystems, Foster City, CA, USA) using GoTaq qPCR Master Mix (Promega) and gene-specific primers for selected phenylpropanoid-, root-associated-, and redox-related genes. The primers used in this study are listed in Supplementary Table 4. Gene expression levels were normalized using *PP2A* as the reference gene. *UBQ10* was also used as an alternative reference gene to evaluate the robustness of expression trends. Relative expression levels were calculated against the mean expression level of the M120 group, which was set to 1 for each gene.

### 2.10 Root hair analysis

Root hair phenotypes were analyzed in mock-treated and SCDY1-treated *Arabidopsis* seedlings grown under 0 or 120 mM NaCl. Five-day-old seedlings grown on 0.5× MS medium with or without 120 mM NaCl were inoculated with mock solution or SCDY1. Root hair phenotypes were analyzed 48 h after inoculation, corresponding to 7-day-old seedlings. The primary root and mature root hair region were observed using a Leica M205 FA stereomicroscope equipped with a Leica DFC7000 T camera (Leica Microsystems, Wetzlar, Germany). Primary root length was measured from root images using ImageJ/Fiji. Root hair zone length was measured as the length of the root hair-bearing region formed after inoculation. The distance from the root apex to the first visible root hair was measured as an index of the position of root hair differentiation. Root hair density and root hair length were quantified in a 3-mm root segment located 2 mm from the primary root tip. Root hair density was calculated as the number of root hairs per millimeter of primary root length. Root hair length was measured for all visible root hairs within the same 3-mm root segment.

### 2.11 Statistical analysis

Statistical analyses and data visualization were performed using R (R Core Team, 2024). For plant growth assays, bacterial trait assays, and root hair phenotyping, statistical tests were selected according to the experimental design. Comparisons between two groups were performed using Student’s t-test. Multiple comparisons among treatment groups were performed using one-way analysis of variance (ANOVA), followed by Tukey’s honestly significant difference (HSD) test. For RT-qPCR analysis, statistical significance between mock-treated and SCDY1-treated seedlings under 120 mM NaCl was assessed using Student’s *t*-test. Differences were considered statistically significant at p < 0.05. For RNA-seq analysis, differential expression was assessed using edgeR based on gene-level read counts, and genes with an absolute log□ fold change of at least 0.585 and an FDR below 0.05 were defined as differentially expressed genes.

## 3 Results

### 3.1 Cube-based screening of quinoa-derived SynComs identifies SynCom DY1 as a candidate salt stress-mitigating consortium

To extend our previous single-isolate analysis of quinoa-associated bacteria (Murata et al., 2026), we designed a cube-based SynCom screening strategy to examine whether salt stress-mitigating activity is primarily associated with individual isolates or with combinations of multiple isolates (Figure 1). A total of 135 bacterial isolates obtained from quinoa seeds and seedlings were used for SynCom construction. Based on partial 16S rRNA gene sequences, the 135 isolates were classified into four bacterial phyla: Bacillota (formerly Firmicutes; 57 isolates), Pseudomonadota (formerly Proteobacteria; 56 isolates), Actinomycetota (formerly Actinobacteria; 16 isolates), and Bacteroidota (formerly Bacteroidetes; 6 isolates) (Figure 1A; Supplementary Table 1). Although the taxonomic composition differed among the five sets, the cube-based design retained broad representation of multiple major plant-associated bacterial phyla across the screening collection. Each 27-isolate set was arranged in a 3 × 3 × 3 cube, and each 3 × 3 layer along the x-, y-, and z-axes was defined as a 9-isolate SynCom (Figure 1B). This design generated nine SynComs per cube and 45 SynComs in total (Figure 1A). An important feature of this design is that each isolate is included in three SynComs, one in each axial direction (Figure 1B). Therefore, activity shared across three SynComs could indicate a dominant isolate-level contribution, whereas layer-specific activity could suggest a combination-dependent effect. This design enabled structured screening of quinoa-derived bacterial communities while retaining traceability of isolate-level and community-level contributions.

**Figure 1.**
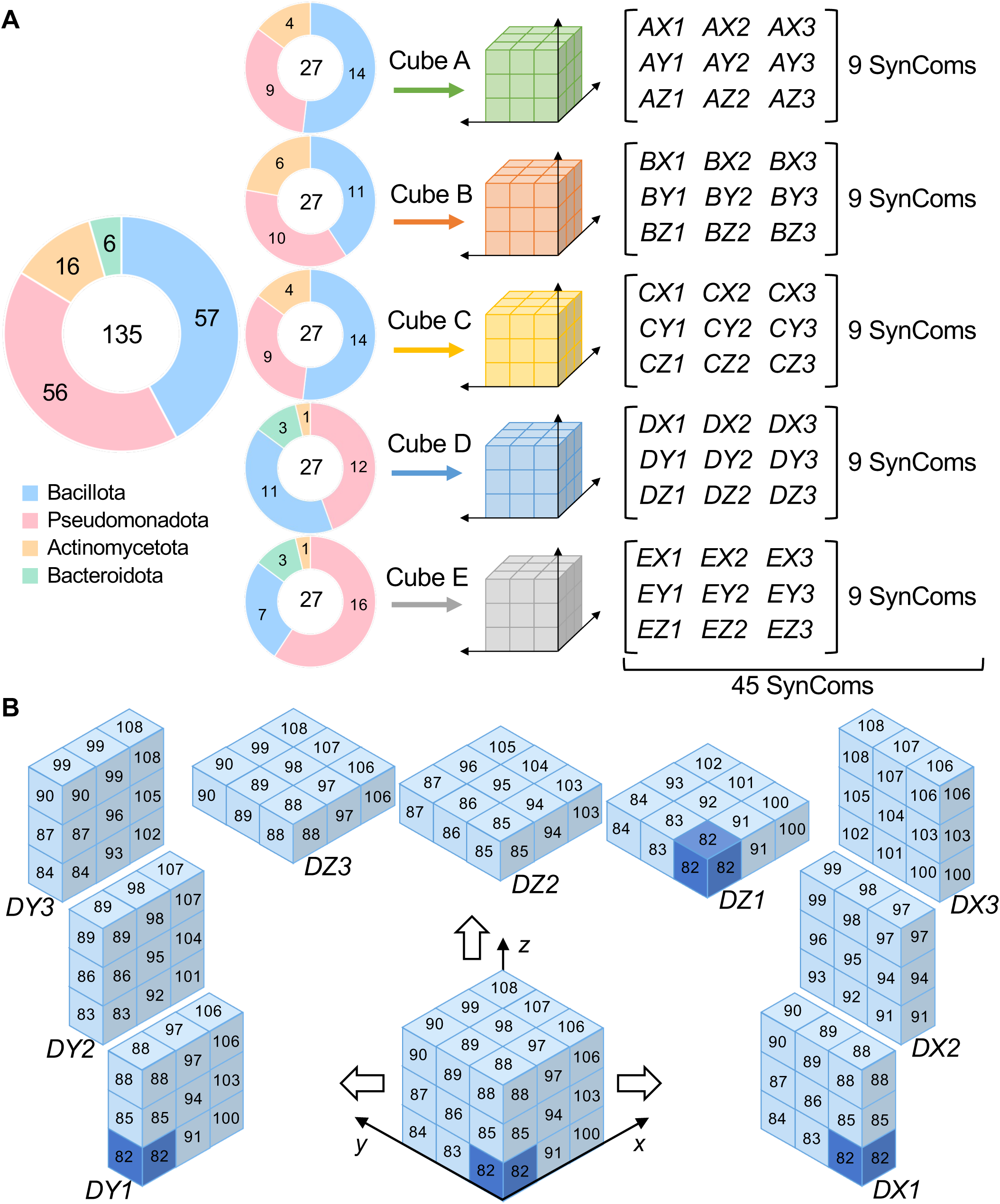
Cube-based screening strategy for identifying quinoa-derived SynComs with salt stress-mitigating activity. (A) Schematic overview of the cube-based SynCom screening design. A total of 135 quinoa-associated bacterial isolates were divided into five independent 27-isolate sets, designated Cubes A–E. The sets were assembled to retain broad taxonomic diversity and to include representatives of major plant-associated bacterial phyla while avoiding overlap among isolates. Each 27-isolate set generated nine 9-isolate SynComs, resulting in 45 SynComs in total. (B) Schematic representation of the layer-based definition of SynComs within a representative 3 × 3 × 3 cube. Each 3 × 3 layer along the x-, y-, and z-axes was defined as one 9-isolate SynCom, so that each isolate was included in three SynComs, one in each axial direction. SynCom DY1 consisted of nine constituent isolates with isolate IDs 82, 85, 88, 91, 94, 97, 100, 103, and 106. If isolate 82 alone were the dominant contributor to the activity, similarly high activity would be expected in the other SynComs containing isolate 82, such as DX1 and DZ1. Instead, the strongest activity was observed in the DY1 layer, supporting the selection of DY1 as a candidate combination-dependent SynCom. SynCom DY1 was selected as SCDY1 for subsequent analyses.

The 45 SynComs were screened for their ability to mitigate salt stress-associated growth inhibition in *Arabidopsis*. The screening was performed under 100 mM NaCl as a primary large-scale bioassay, and SynCom performance was evaluated based on seedling growth relative to mock-treated controls under salinity stress. Among the SynComs tested, SynCom DY1 showed the most consistent positive effect on plant growth under salinity stress and was selected for further analysis as SCDY1 (Supplementary Table 2). SCDY1 was composed of nine predefined quinoa-associated bacterial isolates that had been assigned to the SynCom DY1 layer during the cube-based screening design (Figure 1B). Based on partial 16S rRNA gene sequences, the nine SCDY1 constituent isolates were closely related to *Pseudomonas panipatensis* (isolate 82), *Stenotrophomonas bentonitica* (isolate 85), *Fictibacillus enclensis* (isolate 88), *Novosphingobium subterraneum* (isolate 91), *Herbaspirillum frisingense* (isolate 94), *Acidovorax facilis* (isolate 97), *Bacillus atrophaeus* (isolate 100), *Priestia megaterium* (isolate 103), and *Bacillus subtilis* (isolate 106) (Figure 2A and Supplementary Table 3). These isolates belonged to five bacterial orders, Pseudomonadales, Xanthomonadales, Sphingomonadales, Burkholderiales, and Bacillales, indicating that SCDY1 represented a taxonomically diverse 9-isolate SynCom selected through the cube-based screening strategy.

**Figure 2.**
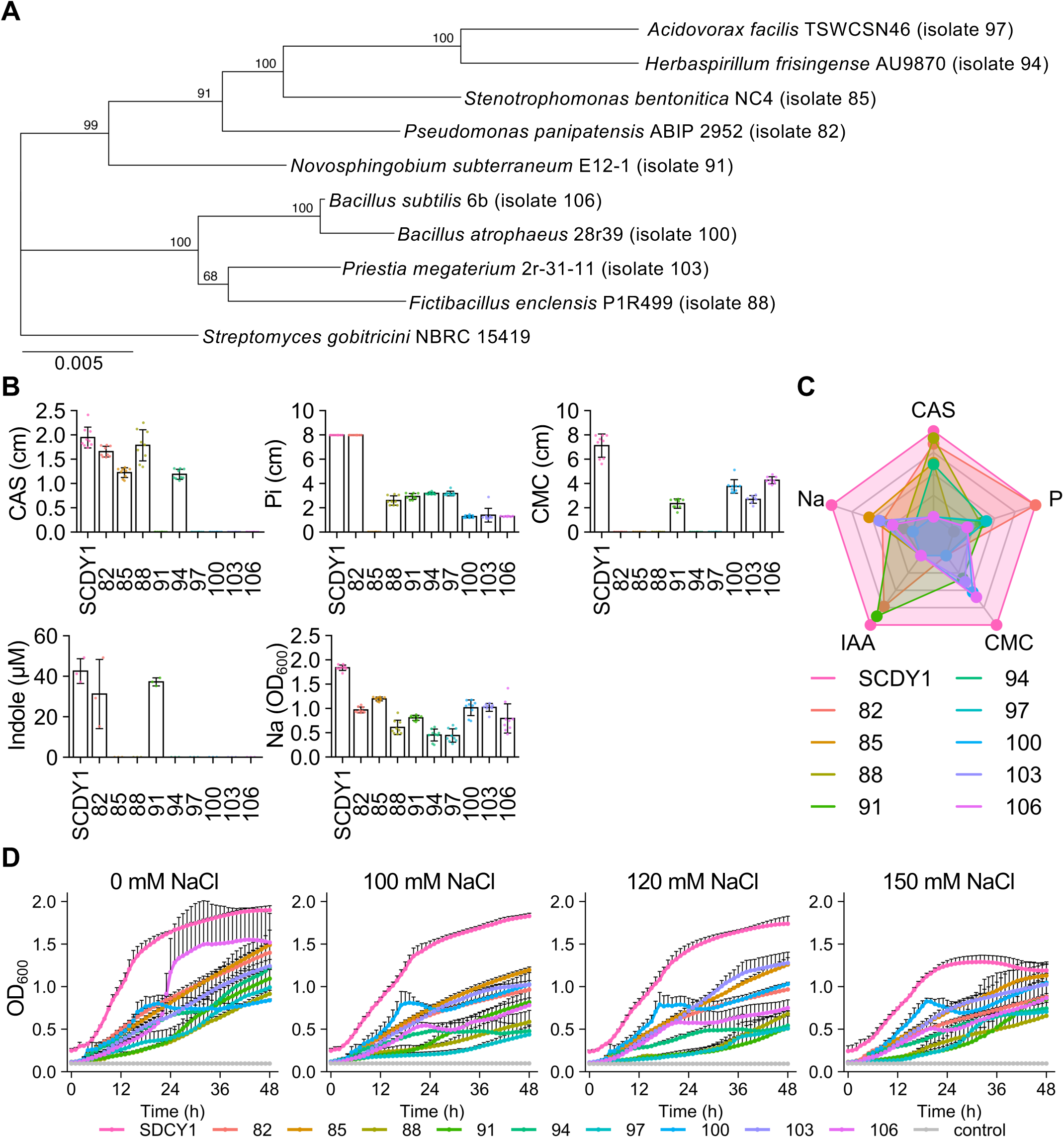
Taxonomic diversity and functional traits of SynCom DY1. (**A**) Phylogenetic tree of the nine SCDY1 member isolates based on partial 16S rRNA gene sequences. The isolates were closely related to *Pseudomonas panipatensis* (isolate 82), *Stenotrophomonas bentonitica* (isolate 85), *Fictibacillus enclensis* (isolate 88), *Novosphingobium subterraneum* (isolate 91), *Herbaspirillum frisingense* (isolate 94), *Acidovorax facilis* (isolate 97), *Bacillus atrophaeus* (isolate 100), *Priestia megaterium* (isolate 103), and *Bacillus subtilis* (isolate 106). *Streptomyces gobitricini* NBRC 15419 was used as an outgroup. Numbers at nodes indicate bootstrap values, and the scale bar indicates substitutions per site. (**B**) Plant growth-promoting traits of SCDY1 and its constituent isolates, including siderophore production, CMC degradation, phosphate solubilization, indole compound production, and growth under saline conditions. Bars represent mean values, and error bars indicate standard deviation. (**C**) Radar chart comparing the multifunctional profiles of SCDY1 and the individual member isolates. (**D**) Growth curves of SCDY1 and individual isolates in media supplemented with 0, 100, 120, or 150 mM NaCl. Bacterial growth was monitored as OD600 over 50 h. Values represent mean ± SD.

### 3.2 SynCom DY1 exhibits a multifunctional profile of plant growth-promoting traits

Having selected SCDY1 through the cube-based screening strategy, we next examined whether this 9-isolate SynCom possessed plant growth-promoting traits that could contribute to salt stress mitigation. We evaluated siderophore production, phosphate solubilization, carboxymethyl cellulose (CMC) degradation, indole compound production, and bacterial growth under saline conditions for SCDY1 and its constituent isolates. The SCDY1 member isolates displayed distinct functional profiles across the tested plant growth-promoting traits (Figure 2B; Supplementary Table 5). Among the individual isolates, isolate 88 showed the highest siderophore production, with a value 3.6% higher than that of SCDY1. Isolate 82 showed the highest phosphate solubilization activity among the individual isolates, reaching a level comparable to that of SCDY1. Isolate 106 showed the highest CMC degradation activity among the individual isolates, corresponding to 59.5% of the SCDY1 value. Isolate 91 exhibited the highest indole compound production among the individual isolates, corresponding to 87.5% of the SCDY1 value. In addition, isolate 85 showed the highest growth under saline conditions among the individual isolates, corresponding to 65.0% of the SCDY1 value. These results indicate that different member isolates of SCDY1 contributed distinct functional capacities rather than uniformly showing the same plant growth-promoting traits. When the multifunctional profiles of the member isolates and SCDY1 were compared by radar chart analysis, SCDY1 showed a broad activity profile across the tested traits (Figure 2C). Except for siderophore production by isolate 88 and phosphate solubilization by isolate 82, SCDY1 exceeded the individual member isolates in all tested plant growth-promoting traits (Figure 2C; Supplementary Table 5). Thus, SCDY1 exhibited a broad multifunctional profile rather than a single dominant plant growth-promoting trait, consistent with the distribution of distinct functional capacities among its member isolates.

We further examined bacterial growth under saline conditions by monitoring optical density over 50 h in media containing 0, 100, 120, or 150 mM NaCl. Across all NaCl concentrations tested, SCDY1 showed faster growth than each of its individual constituent isolates (Figure 2D). Together, these results show that SCDY1 combines multifunctional plant growth-promoting traits with robust growth under saline conditions.

### 3.3 SynCom DY1 promotes *Arabidopsis* growth in a salinity-dependent manner

To determine whether the selected consortium could mitigate salt stress-associated growth inhibition in plants, we examined the effects of SCDY1 inoculation on *Arabidopsis* seedlings grown under non-saline and saline conditions. *Arabidopsis* was used as a sensitive model host to quantitatively evaluate root growth and biomass responses to microbial inoculation under controlled salinity stress. Seedlings were transferred to medium with or without NaCl and inoculated with SCDY1 or mock solution near the root tip region. Under non-saline conditions, primary root growth was faster in mock-treated seedlings than in SCDY1-treated seedlings until the roots reached the bottom of the plate at 187 h after inoculation (Figure 3A, B). In contrast, fresh and dry weights did not differ significantly between mock- and SCDY1-treated seedlings under non-saline conditions (Figure 3C, D). These results indicate that SCDY1-mediated growth promotion was salinity-dependent rather than constitutive. Under 100 mM NaCl conditions, SCDY1-treated seedlings showed enhanced root growth compared with mock-treated controls until the roots reached the bottom of the plate at 306 h after inoculation (Figure 3A, E). Quantitative analysis showed that SCDY1 significantly promoted primary root elongation under salt stress. Consistent with this root growth phenotype, fresh and dry weights were significantly higher in SCDY1-treated seedlings than in mock-treated seedlings under 100 mM NaCl conditions (Figure 3F, G). These results indicate that the growth-promoting activity of SCDY1 is more evident under salinity stress than under non-stress conditions.

**Figure 3.**
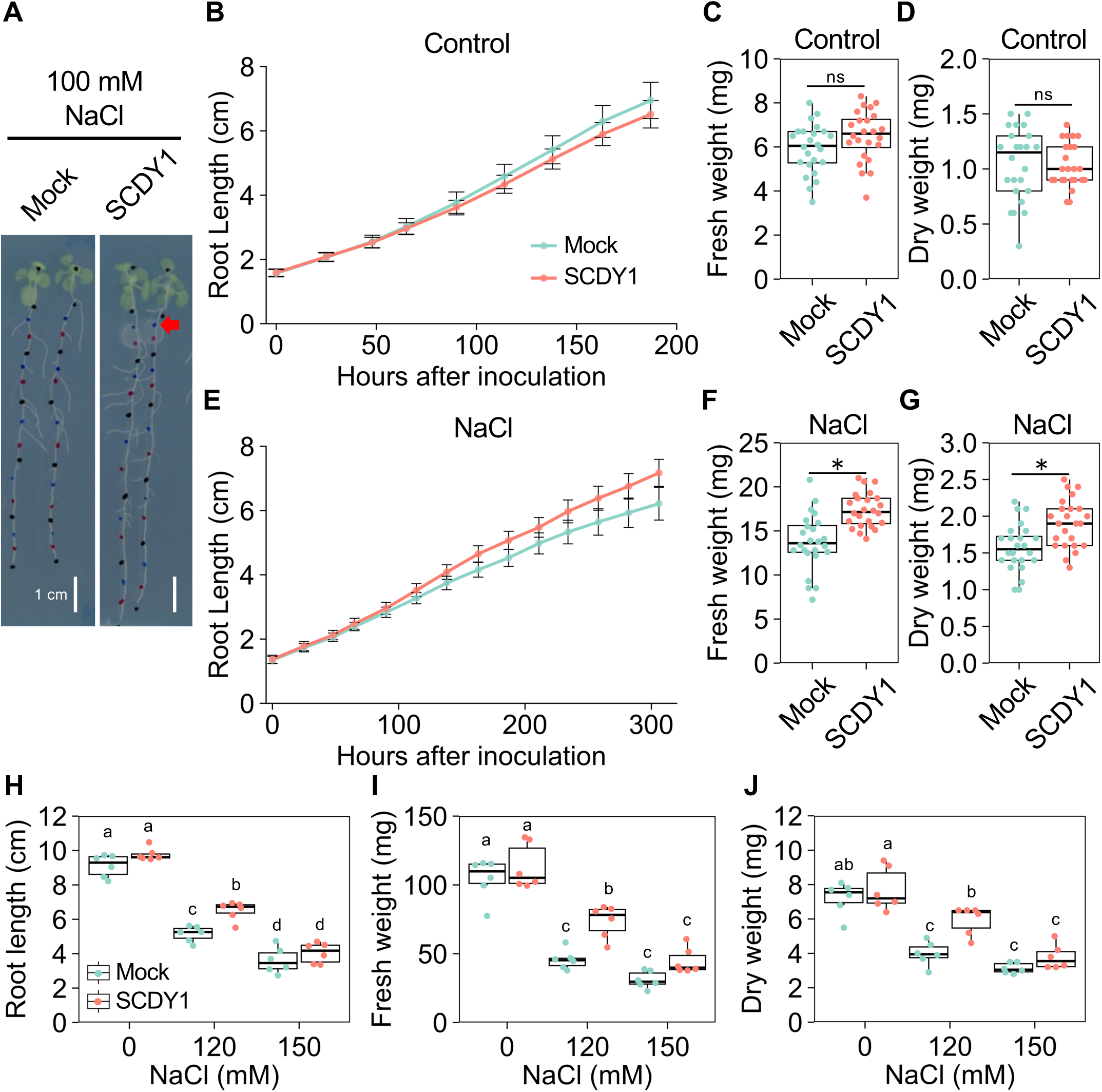
Salinity-dependent growth promotion of *Arabidopsis* seedlings by SCDY1. (A) Representative images of mock-treated and SCDY1-treated *Arabidopsis* seedlings grown under 100 mM NaCl. Arrows indicate the inoculation points of mock solution or SCDY1 near the root tip region. Colored marks indicate sequential root tip positions recorded during time-course measurement of primary root elongation. Scale bars = 1 cm. (B, E) Time-course analysis of primary root elongation under non-saline control conditions (B) and 100 mM NaCl (E). (C, D, F, G) Fresh weight (C, F) and dry weight (D, G) of mock-treated and SCDY1-treated seedlings under non-saline control conditions (C, D) and 100 mM NaCl (F, G). (H–J) Primary root length (H), fresh weight (I), and dry weight (J) of mock-treated and SCDY1-treated seedlings grown under 0, 120, or 150 mM NaCl. SCDY1 inoculation significantly increased root length and biomass under 120 mM NaCl but not under 0 or 150 mM NaCl. Line graphs show mean ± SD. Box plots show medians, interquartile ranges, whiskers, and individual biological replicates. Asterisks and different lowercase letters indicate significant differences at *p* < 0.05, as determined using Student’s *t*-test and Tukey’s HSD test.

To further evaluate the salinity range in which SCDY1 exerts its growth-promoting effect, *Arabidopsis* seedlings were exposed to 0, 120, or 150 mM NaCl. SCDY1 inoculation significantly increased primary root length, fresh weight, and dry weight under 120 mM NaCl, whereas this effect was not detected under 0 mM or 150 mM NaCl (Figure 3H–J). These results suggest that SCDY1 is particularly effective under moderate salinity stress, where plant growth is inhibited but not severely suppressed. Because the SCDY1-dependent growth phenotype was most clearly observed at 120 mM NaCl, this condition was selected for subsequent transcriptome analysis to investigate host responses associated with SCDY1-mediated salt stress mitigation.

To confirm that SCDY1-derived bacteria were present in the plant assay system after inoculation, we performed a culture-dependent re-isolation assay. Seedlings inoculated with SCDY1 or mock solution were homogenized after growth under salinity stress, and the homogenates were plated on bacterial culture media. Bacterial colonies were recovered from SCDY1-inoculated seedlings on multiple media, whereas no colonies were obtained from mock-treated seedlings. Colony PCR followed by sequence analysis identified colonies corresponding to several SCDY1 constituent isolates, including isolates 82, 85, 94, and 106 (Supplementary Table 6). These results indicate that at least a subset of SCDY1 constituent bacteria was recoverable from inoculated seedlings under salinity stress conditions.

### 3.4 SCDY1 modulates the *Arabidopsis* salt stress transcriptome under moderate salinity stress

Because SCDY1 promoted *Arabidopsis* growth most clearly under 120 mM NaCl in our phenotypic assay, this condition was used as a moderate salinity stress condition for transcriptome analysis. Total RNA was extracted from seedlings at 100 h after inoculation, when SCDY1-dependent differences in growth phenotype were clearly detectable under 120 mM NaCl. Four treatment groups were analyzed: mock-treated seedlings under non-saline conditions (M0), mock-treated seedlings under 120 mM NaCl (M120), SCDY1-treated seedlings under non-saline conditions (S0), and SCDY1-treated seedlings under 120 mM NaCl (S120). Differentially expressed genes (DEGs) were identified using four pairwise comparisons: M120 vs M0, S120 vs S0, S0 vs M0, and S120 vs M120. Salt stress caused the largest transcriptional changes in both mock- and SCDY1-treated plants, with 2,256 DEGs in M120 vs M0 and 1,778 DEGs in S120 vs S0 (Figure 4A; Supplementary Table 7). Direct comparisons between SCDY1- and mock-treated plants identified fewer DEGs, with 341 DEGs under non-saline conditions and 567 DEGs under 120 mM NaCl conditions. The direction of SCDY1-responsive transcriptional changes differed between non-saline and saline conditions. Under non-saline conditions, most SCDY1-responsive DEGs were upregulated, with 299 upregulated and 42 downregulated genes in S0 vs M0. In contrast, under 120 mM NaCl, SCDY1-responsive DEGs included similar numbers of upregulated and downregulated genes, with 256 upregulated and 311 downregulated genes in S120 vs M120. These results indicate that salt stress was the major driver of transcriptional variation, whereas SCDY1 induced a distinct transcriptional response that was smaller than the salinity-driven response. The balanced up- and downregulation observed under 120 mM NaCl suggests that SCDY1 modulated, rather than simply activated, the host salt stress response. Volcano plot analysis further showed that many SCDY1-responsive DEGs displayed moderate fold changes rather than large expression shifts (Figure 4B). Principal component analysis (PCA) showed that PC1, which explained 45.6% of the total variance, separated samples primarily according to salinity treatment, whereas PC2, which explained 10.9% of the variance, reflected variation associated with SCDY1 treatment (Figure 4C). SCDY1- and mock-treated samples showed limited separation under non-saline conditions but were more clearly separated under 120 mM NaCl, and biological replicates clustered closely within each treatment group.

**Figure 4.**
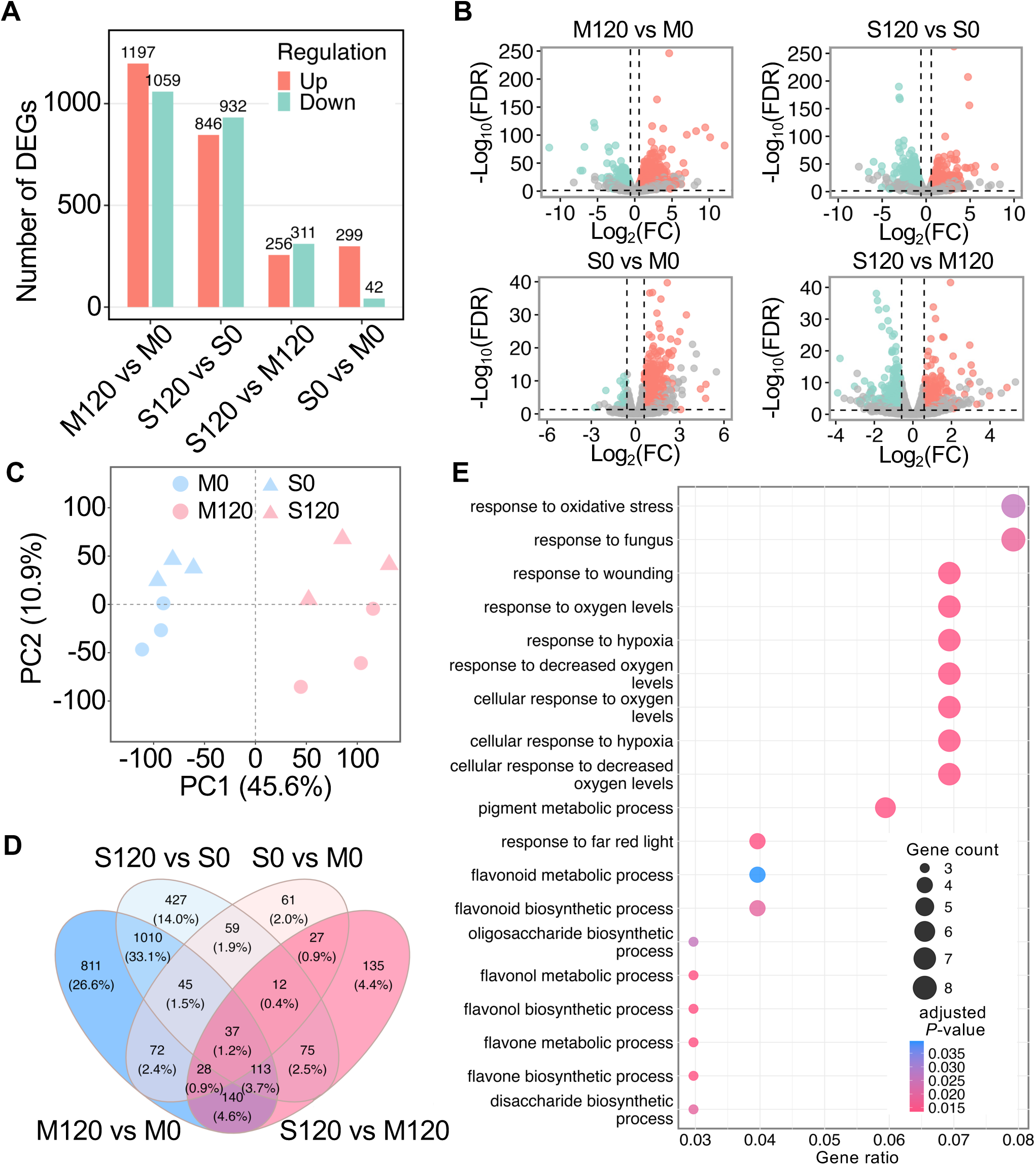
Transcriptome analysis of SCDY1-treated *Arabidopsis* seedlings under moderate salinity stress. (A) Numbers of differentially expressed genes (DEGs) identified in four pairwise comparisons: M120 vs M0, S120 vs S0, S120 vs M120, and S0 vs M0. Upregulated and downregulated DEGs are shown separately. (B) Volcano plots of DEGs identified in the four pairwise comparisons. Red and blue points indicate upregulated and downregulated genes, respectively, and gray points indicate genes that were not significantly differentially expressed. (C) Principal component analysis (PCA) of RNA-seq samples from the four treatment groups. PC1 explained 45.6% of the total variance and primarily separated samples according to salinity treatment, whereas PC2 explained 10.9% of the variance and reflected variation associated with SCDY1 treatment. (D) Venn diagram showing the overlap of DEGs among the four pairwise comparisons. A total of 3,052 unique DEGs were detected across all comparisons, and 135 DEGs were uniquely identified in the S120 vs M120 comparison. (E) Gene Ontology enrichment analysis of the 135 SCDY1-specific DEGs uniquely identified in the S120 vs M120 comparison. Enriched biological process terms included stress-associated responses such as response to oxidative stress, response to fungus, and response to wounding. Dot size indicates the number of genes assigned to each GO term, and dot color indicates the adjusted *p* value. M0, mock-treated seedlings under non-saline conditions; M120, mock-treated seedlings under 120 mM NaCl; S0, SCDY1-treated seedlings under non-saline conditions; S120, SCDY1-treated seedlings under 120 mM NaCl.

To further examine the specificity of SCDY1-responsive transcriptional changes, we compared the overlap of DEGs among the four pairwise comparisons. A total of 3,052 unique DEGs were detected across all comparisons. Among these, 135 DEGs were uniquely identified in the S120 vs M120 comparison and were therefore designated SCDY1-specific DEGs under salinity stress (Figure 4D; Supplementary Table 8). Gene Ontology enrichment analysis of these SCDY1-specific DEGs indicated enrichment of processes related to oxidative stress responses, responses to fungus, and responses to wounding (Figure 4E; Supplementary Table 9). Together, these results indicate that SCDY1 modulates the *Arabidopsis* transcriptome in a salinity-dependent manner, with distinct stress-responsive gene sets detected under 120 mM NaCl.

### 3.5 SCDY1-responsive gene clusters reveal stress- and root-associated transcriptional modules under salinity stress

To further characterize the biological processes associated with SCDY1-dependent transcriptional changes under salt stress, we performed hierarchical clustering of DEGs identified across the treatment comparisons. The DEGs were classified into 10 clusters according to their expression patterns across the four treatment groups (Figure 5A; Supplementary Table 10). These clusters revealed distinct transcriptional modules associated with salinity stress and SCDY1 inoculation, suggesting that SCDY1 modulates specific subsets of stress-responsive genes rather than globally altering host gene expression. Among the 10 clusters, Cluster 1 contained a major group of genes enriched for biological processes related to response to water deprivation, response to oxidative stress, and response to jasmonic acid (Figure 5B). Cluster 3 was enriched for genes associated with response to oxygen levels, cellular response to oxygen levels, and response to hypoxia (Figure 5C). Cluster 7 contained a smaller set of genes enriched for plant-type cell wall organization or biogenesis, plant epidermis development, and root epidermal cell differentiation (Figure 5D). These clusters linked SCDY1-responsive transcriptional changes under salinity stress to water stress, oxygen/redox regulation, and root cell wall- or epidermis-related processes.

**Figure 5.**
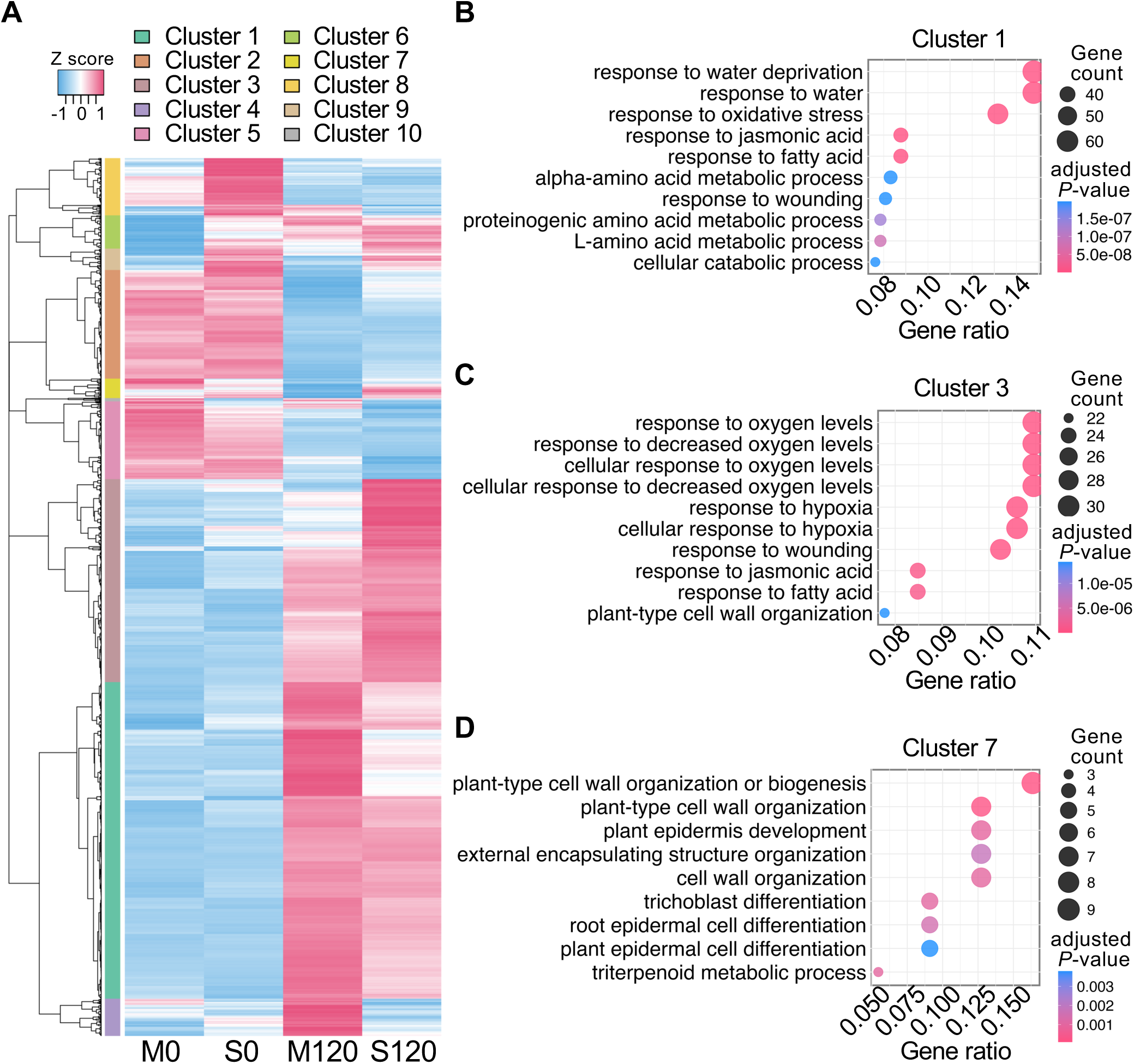
SCDY1-responsive transcriptional modules associated with stress- and root-related processes under salinity stress. (**A**) Hierarchical clustering of DEGs across M0, S0, M120, and S120. DEGs were classified into 10 clusters based on their expression patterns. Red and blue indicate relatively high and low expression levels, respectively. (B–D) Gene Ontology enrichment analysis of representative clusters. Cluster 1 was enriched for water deprivation-, oxidative stress-, and jasmonic acid-related responses (B). Cluster 3 was enriched for oxygen level-, hypoxia-, and wounding-related responses (C). Cluster 7 was enriched for plant-type cell wall organization or biogenesis, plant epidermis development, and root epidermal cell differentiation (D). Dot size indicates gene count, and dot color indicates the adjusted *P* value. M0, mock-treated seedlings under non-saline conditions; S0, SCDY1-treated seedlings under non-saline conditions; M120, mock-treated seedlings under 120 mM NaCl; S120, SCDY1-treated seedlings under 120 mM NaCl.

We next performed Gene Ontology enrichment analysis of SCDY1-responsive DEGs to obtain a broader overview of enriched biological functions. In the biological process category, enriched terms included response to water, response to oxidative stress, and response to oxygen levels (Figure 6A; Supplementary Table 11). In the molecular function category, enriched terms included monooxygenase activity, oxidoreductase activity, antioxidant activity, and glucosyltransferase activity (Figure 6B; Supplementary Table 12). Together, these clustering and GO enrichment analyses associated SCDY1-responsive genes with stress-responsive, redox-related, and root cell wall-associated processes in *Arabidopsis*.

**Figure 6.**
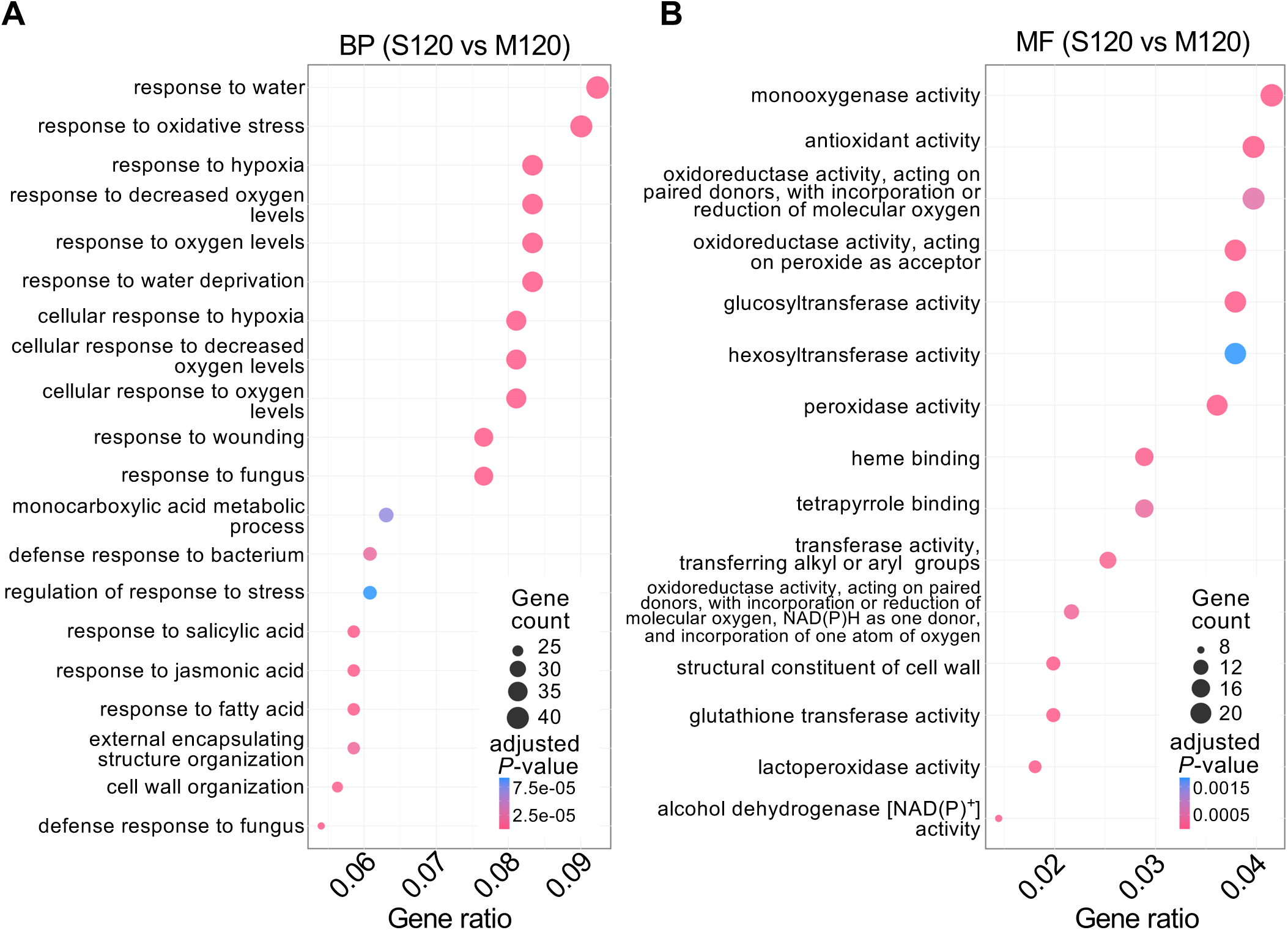
Gene Ontology enrichment analysis of SCDY1-responsive DEGs under salinity stress. (A, B) Dot plots showing enriched Gene Ontology terms among SCDY1-responsive DEGs identified in the S120 vs M120 comparison. Biological process terms are shown in (A), and molecular function terms are shown in (B). Enriched biological process terms included response to water, response to oxidative stress, and response to hypoxia, whereas enriched molecular function terms included monooxygenase activity, antioxidant activity, and oxidoreductase activity. Dot size indicates the number of genes assigned to each GO term, and dot color indicates the adjusted *P* value. S120, SCDY1-treated seedlings under 120 mM NaCl; M120, mock-treated seedlings under 120 mM NaCl.

### 3.6 SCDY1-responsive genes are associated with phenylpropanoid biosynthesis, root-associated responses, and glutathione metabolism

To identify metabolic and signaling pathways associated with SCDY1-responsive transcriptional changes under salt stress, we performed KEGG pathway enrichment analysis using DEGs identified in the S120 vs M120 comparison. KEGG analysis showed enrichment of genes associated with phenylpropanoid biosynthesis, glutathione metabolism, and plant MAPK signaling (Figure 7; Supplementary Table 13). Among these pathways, phenylpropanoid biosynthesis and glutathione metabolism were most directly related to the cell wall-associated and redox-related responses identified by GO enrichment analysis.

**Figure 7.**
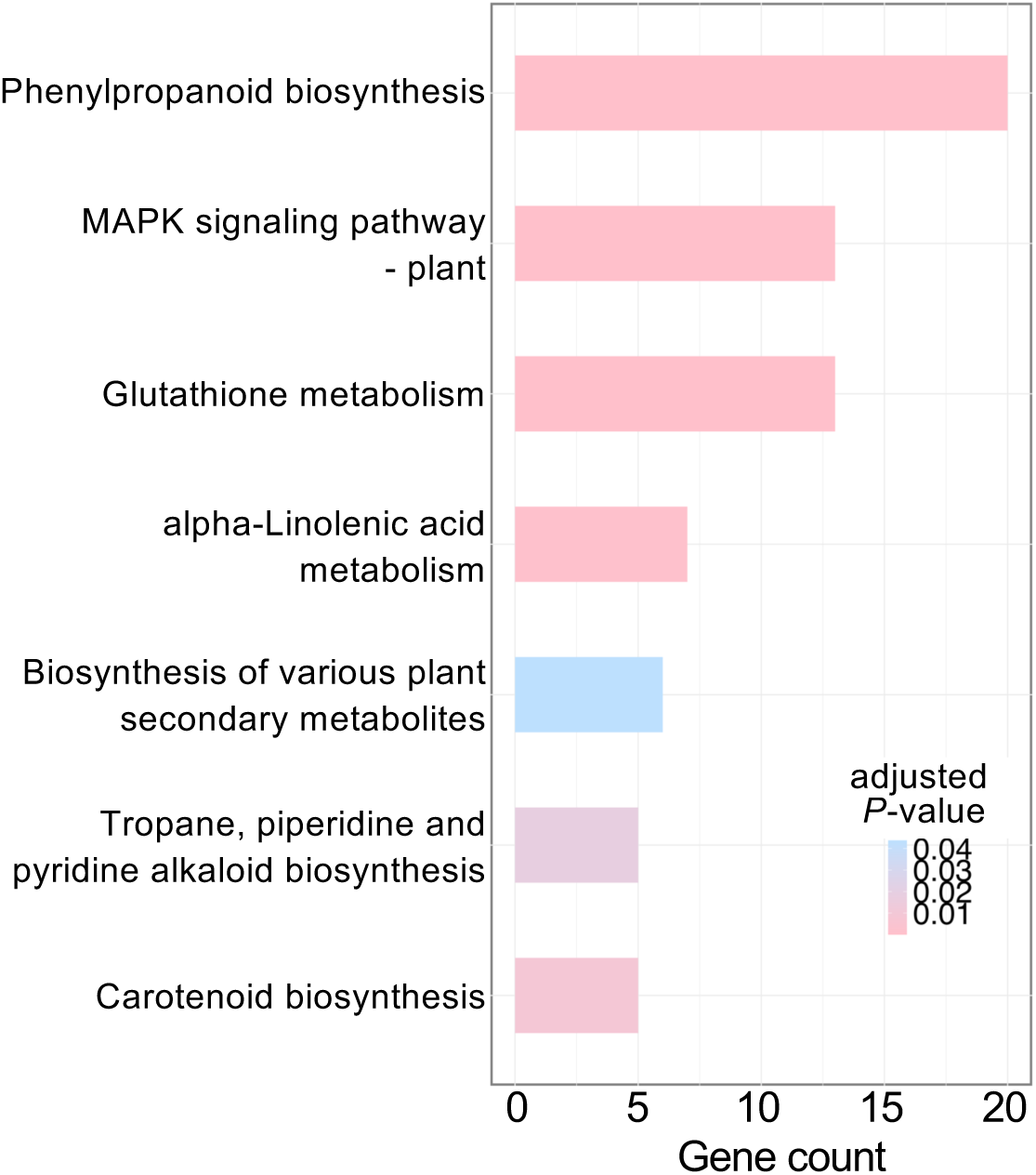
KEGG pathway enrichment analysis of SCDY1-responsive DEGs under salinity stress. (A) KEGG pathway enrichment analysis of differentially expressed genes identified in the S120 vs M120 comparison. Bars indicate the number of DEGs assigned to each enriched KEGG pathway, and color indicates the adjusted *P* value. Phenylpropanoid biosynthesis, plant MAPK signaling, and glutathione metabolism were among the enriched pathways associated with SCDY1 treatment under 120 mM NaCl. (B) Heatmap representation of DEGs assigned to the phenylpropanoid biosynthesis and glutathione metabolism pathways. Gene expression changes are shown as log2 fold change values in the S120 vs M120 comparison. Red indicates upregulation in SCDY1-treated seedlings, whereas blue indicates downregulation. S120, SCDY1-treated seedlings under 120 mM NaCl; M120, mock-treated seedlings under 120 mM NaCl.

Phenylpropanoid biosynthesis showed a prominent transcriptional signature, with 20 assigned DEGs, including 12 upregulated and 8 downregulated genes in SCDY1-treated plants under salt stress (Figure 7; Supplementary Table 14). Several class III peroxidase genes, including *AtPRX1*, *PRX2*, *PER7*, *PRX8*, *PRX35*, *PRX37*, *PER39*, *PRX44*, and *PRX73*, were upregulated in SCDY1-treated seedlings. Root hair-specific genes, including *RHS18* and *RHS19*, were also upregulated (Supplementary Table 14). Thus, the phenylpropanoid-related DEG set included genes associated with cell wall modification and root epidermal responses.

Genes assigned to glutathione metabolism showed a predominantly downregulated pattern in SCDY1-treated plants under salt stress. Thirteen DEGs were assigned to this pathway, including glutathione S-transferase genes such as *GSTU1*, *GSTU4*, *GSTU7*, *GSTU17*, *GSTU22*, *GSTU24*, *GSTU25*, *GSTF2*, and *GSTF6*, as well as redox-related genes including *DHAR2* and *G6PD3* (Figure 7; Supplementary Table 15). Stress-responsive genes such as *ERD9* and *ERD11* also showed reduced expression (Supplementary Table 15), linking SCDY1 treatment to glutathione- and redox-associated transcriptional responses under salinity stress.

We next performed targeted RT-qPCR analysis of selected genes using the same RNA samples used for RNA-seq analysis. The selected genes included class III peroxidase genes (*PRX44*, *PER39*, *PRX2*, and *PRX35*), root hair- or root epidermis-related genes (*RHS18* and *RHS19*), and glutathione metabolism- or redox-related genes (*GSTF6*, *GSTU24*, *GSTU7*, *DHAR2*, and *ERD9*) (Figure 8). These genes were selected based on the RNA-seq and pathway analyses (Supplementary Tables 14 and 15). RT-qPCR analysis supported the RNA-seq expression patterns: *PRX44*, *PER39*, *PRX2*, *PRX35*, *RHS18*, and *RHS19* showed increased expression in SCDY1-treated seedlings, whereas *GSTU24*, *GSTU7*, and *DHAR2* showed reduced expression (Figure 8). Together, KEGG pathway analysis and targeted RT-qPCR analysis linked SCDY1-responsive transcriptional changes to phenylpropanoid biosynthesis, root hair-related responses, and glutathione-associated redox metabolism under salinity stress.

**Figure 8.**
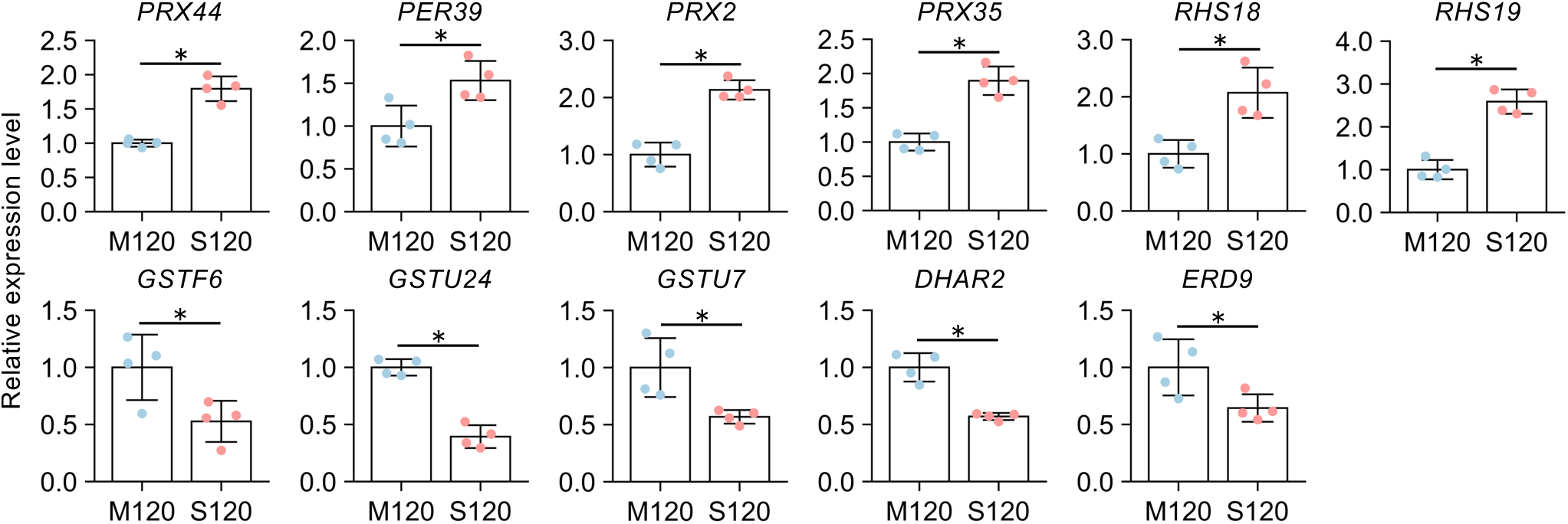
Targeted RT-qPCR analysis of selected SCDY1-responsive genes under salinity stress. Relative expression levels of selected phenylpropanoid- and cell wall-related genes (PRX44, PER39, PRX2, and PRX35), root hair- or root epidermis-related genes (RHS18 and RHS19), and glutathione metabolism- or redox-related genes (GSTF6, GSTU24, GSTU7, DHAR2, and ERD9) were analyzed in mock-treated and SCDY1-treated Arabidopsis seedlings grown under 120 mM NaCl. Gene expression levels were quantified by RT-qPCR and normalized to PP2A as the reference gene. Expression levels are shown as fold changes relative to the mean expression level of mock-treated seedlings under 120 mM NaCl (M120), which was set to 1 for each gene. Bars represent means ± SD, and individual points represent biological replicates (*n* = 4). Statistical significance between M120 and S120 was assessed using a two-sided Student’s t-test. Asterisks indicate significant differences (\**p* < 0.05). M120, mock-treated seedlings under 120 mM NaCl; S120, SCDY1-treated seedlings under 120 mM NaCl.

### 3.7 SCDY1 treatment enhances root hair-related phenotypes in *Arabidopsis* seedlings

Because transcriptome and targeted RT-qPCR analyses linked SCDY1 treatment to root epidermis- and root hair-related genes, we examined root hair phenotypes under non-saline and saline conditions. Five-day-old seedlings grown under 0 or 120 mM NaCl were inoculated with mock solution or SCDY1, and primary root and root hair phenotypes were analyzed 48 h after inoculation (Figure 9A). Primary root length was reduced under 120 mM NaCl but did not differ significantly between mock- and SCDY1-treated seedlings under either NaCl condition (Figure 9B), allowing root hair-related phenotypes to be evaluated without detectable treatment-dependent differences in primary root elongation.

**Figure 9.**
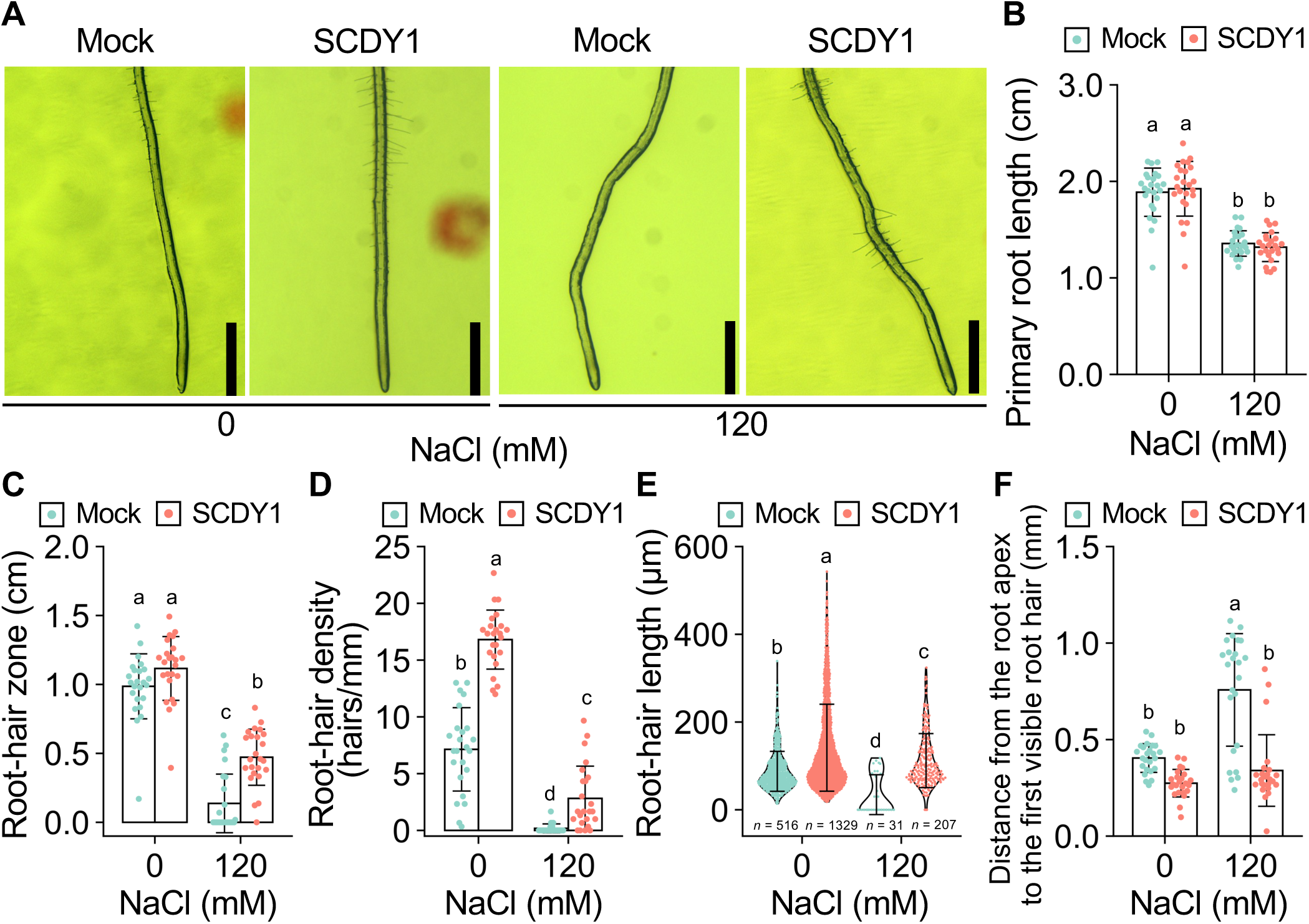
SCDY1 treatment enhances root hair-related phenotypes in *Arabidopsis* seedlings. Five-day-old *Arabidopsis* seedlings grown on half-strength MS medium with or without 120 mM NaCl were inoculated with mock solution or SCDY1, and primary root and root hair phenotypes were analyzed 48 h after inoculation. (A) Representative images of root hairs in mock-treated and SCDY1-treated seedlings under 0 and 120 mM NaCl conditions. Scale bar = 1 mm. (B) Primary root length. (C) Root hair zone length, measured as the maturation zone formed after inoculation to the last visible root hair. (D) Root hair density, quantified in a 3-mm root segment located 2 mm from the primary root tip and expressed as the number of root hairs per millimeter of primary root length. (E) Root hair length measured in the same root region. (F) Distance from the root apex to the first visible root hair, used as an index of the position of root hair differentiation. For panels B–D, bars represent mean ± SD, and dots indicate individual biological replicates (*n* = 24). For panel E, violin plots show the distribution of root hair length, box plots indicate the median and interquartile range, and dots indicate individual root hair measurements. Sample sizes for panel E are indicated below each violin plot. Different letters indicate significant differences among treatment groups based on one-way ANOVA followed by Tukey’s HSD test (p < 0.05).

Root hair zone length did not differ significantly between mock- and SCDY1-treated seedlings under 0 mM NaCl, but was significantly greater in SCDY1-treated seedlings than in mock-treated seedlings under 120 mM NaCl (Figure 9C). Root hair density and root hair length were significantly higher in SCDY1-treated seedlings than in mock-treated seedlings under both 0 and 120 mM NaCl conditions (Figure 9D, E). Although salinity reduced root hair density and root hair length, SCDY1-treated seedlings maintained higher values than mock-treated seedlings under salinity stress. In addition, the distance from the root apex to the first visible root hair was shorter in SCDY1-treated seedlings than in mock-treated seedlings under both NaCl conditions (Figure 9F), suggesting that SCDY1 treatment is associated with an earlier onset of root hair differentiation. Together, these results show that SCDY1 enhances root hair-related phenotypes and shifts visible root hair formation closer to the root apex without detectable treatment-dependent changes in primary root length, providing phenotypic support for SCDY1-associated root epidermal responses under moderate salinity stress.

## 4 Discussion

Here, we developed a cube-based SynCom screening strategy to identify quinoa-derived bacterial consortia that mitigate salt stress-associated growth inhibition in plants. By screening 135 quinoa-associated bacterial isolates as 45 defined 9-isolate SynComs, we identified SynCom DY1 (SCDY1) as a taxonomically diverse consortium that promoted *Arabidopsis* growth under moderate salinity stress (Figures 1, 3). A key feature of this screening design is that it preserves structured combinatorial diversity while keeping the number of SynComs within a practical screening scale (Figure 1). This allowed us to explore different isolate combinations while providing initial information on whether beneficial activity was more consistent with isolate-level or combination-dependent effects. SCDY1 exhibited a broad multifunctional profile, including traits related to nutrient mobilization, indole compound production, iron acquisition, cell wall polymer degradation, and bacterial growth under saline conditions (Figure 2). Because these traits were distributed among different member isolates, SCDY1 appears to integrate complementary functional capacities that may contribute to its ability to mitigate plant growth inhibition under salt stress.

SCDY1 promoted *Arabidopsis* growth in a salinity-dependent manner, with the clearest effects observed under 100–120 mM NaCl (Figure 3). This pattern suggests that SCDY1 functions most effectively under moderate salinity stress, where plant growth is constrained but remains responsive to microbial modulation. The absence of a clear growth-promoting effect under non-saline or more severe salinity conditions further supports the view that SCDY1 acts as a stress-context-dependent consortium rather than as a constitutive growth stimulant. Bacterial re-isolation from inoculated seedlings showed that several SCDY1 constituent bacteria were recoverable during the assay period. The assay did not resolve the persistence, abundance, or spatial distribution of all consortium members, but it confirmed the presence of recoverable SCDY1-derived bacteria under the conditions in which growth promotion was observed. Together, these findings support a model in which SCDY1 promotes plant growth most effectively under moderate salinity stress, while maintaining recoverable bacterial members in the plant assay system.

Host transcriptome analysis provided insight into plant responses associated with SCDY1-mediated growth promotion under salt stress. Salt stress is known to restrict root growth by affecting root meristem activity, auxin distribution, redox status, water relations, and cell expansion (Jiang et al., 2016; Liu et al., 2015). In our dataset, salinity was the dominant driver of transcriptional variation, whereas SCDY1 induced a distinct transcriptional response that was smaller than the salinity-driven response (Figure 4). The balanced up- and downregulation of SCDY1-responsive genes under 120 mM NaCl suggests that SCDY1 did not simply activate a general stress response but instead modulated specific components of the host salt stress program. GO, clustering, KEGG, and RT-qPCR analyses consistently associated SCDY1 treatment with stress-responsive, redox-related, phenylpropanoid-associated, glutathione-associated, and root epidermis-related transcriptional changes (Figures 5–8). Among these responses, the enrichment of cell wall organization, plant epidermis development, and root epidermal cell differentiation terms was particularly relevant to the root hair-related phenotypes observed in SCDY1-treated seedlings, because root epidermal cells and root hairs are highly plastic structures that respond to nutrient availability, water status, and abiotic stress (Zhang et al., 2023). Root hair phenotyping further showed that SCDY1 enhanced multiple root hair-related traits and shifted visible root hair formation closer to the root apex, without detectable treatment-dependent changes in primary root length at the analyzed time point (Figure 9). Although the causal relationship between root epidermal responses and growth promotion remains to be tested, the induction of class III peroxidase genes and root hair-related genes provides candidate molecular links between SCDY1 treatment, cell wall-or redox-associated processes, and root epidermal responses (Figures 8, 9). Plant class III peroxidases participate in ROS-associated reactions, cell wall modification, lignification, and cell expansion, and some members have been implicated in root hair growth through apoplastic ROS and cell wall regulation (Cosio and Dunand, 2009; Marjamaa et al., 2009; Pacheco et al., 2022; Passardi et al., 2004). Together, this study provides a practical framework for moving from large plant-associated bacterial collections to defined SynComs with testable functions in plant stress tolerance.

Previous stress-microbiome and SynCom studies provide an important context for the present work. A halophyte-associated microbiome was previously shown to help non-host plants withstand salinity, supporting the idea that stress-adapted plants can harbor microbial resources that improve plant stress performance (Yuan et al., 2016). Similarly, a synthetic bacterial community derived from a desert rhizosphere was shown to confer salt stress resilience to tomato even in the presence of a soil microbiome (Schmitz et al., 2022). In line with these studies, our results support the view that plants adapted to stressful environments can serve as useful sources of beneficial microorganisms. The distinct contribution of the present work is the cube-based screening framework, which enabled evaluation of a large quinoa-derived isolate collection at a practical scale while preserving the traceability of isolate combinations (Figure 1). The present study establishes a controlled screening platform that can now be extended in several directions. Dropout SynCom analysis, reduced SynCom reconstruction, and comparison with selected individual isolates will help define the member isolates or functional groups responsible for SCDY1 activity. Isolate-specific qPCR, amplicon sequencing, or fluorescent labeling will clarify the persistence, spatial distribution, and relative abundance of SCDY1 members on or near plant roots. Further analyses of cell wall modification, ROS distribution, and root hair-related pathways will help test how root epidermal responses contribute to growth promotion. Finally, testing SCDY1 or related quinoa-derived SynComs in quinoa, crop species, and soil-based systems will be important for evaluating the broader applicability of this approach. Thus, the cube-based SynCom strategy developed here provides a practical starting point for moving from large stress-adapted bacterial collections toward defined microbial communities with testable functions in plant stress performance.

## Conclusions

This study developed a cube-based screening strategy for identifying beneficial SynComs from quinoa-associated bacterial collections. Using this approach, we identified SCDY1 as a taxonomically diverse 9-isolate SynCom with multifunctional plant growth-promoting traits and salinity-dependent growth-promoting activity in *Arabidopsis*. Bacterial re-isolation supported the presence of SCDY1-derived bacteria in the plant assay system. Host transcriptome analysis, targeted RT-qPCR analysis, and root hair phenotyping linked SCDY1 treatment to stress-responsive and redox-associated transcriptional changes as well as enhanced root hair-related epidermal phenotypes, including a shift in visible root hair formation toward the root apex, under moderate salinity stress. These findings suggest that quinoa-associated bacterial communities can enhance plant performance under salinity stress through complementary microbial traits and modulation of host root-associated stress responses. Together, this study provides a practical framework for moving from large plant-associated bacterial collections to defined SynComs with testable functions in plant performance under salinity stress.

## Data availability statement

The datasets generated in this study are available at the INSDC (DDBJ, EMBL, and GenBank) under BioProject accession PRJDB42650 (https://ddbj.nig.ac.jp/search/entry/bioproject/PRJDB42650/).

## Author contributions

HD: Investigation, Data curation, Formal analysis, Methodology, Validation, Visualization, Writing – original draft, Writing – review & editing. YM: Conceptualization, Formal analysis, Investigation, Methodology, Resources, Supervision, Writing – original draft, Writing – review & editing. YK: Formal analysis, Investigation, Data curation, Supervision, Validation, Visualization, Methodology, Writing – original draft, Writing – review & editing. SN: Investigation, Data curation, Formal analysis, Visualization, Methsodology, Writing – original draft, Writing – review & editing. TO: Methodology, Project administration, Supervision, Validation, Visualization, Writing – original draft, Writing – review & editing. YF: Conceptualization, Methodology, Funding acquisition, Project administration, Resources, Supervision, Validation, Visualization, Writing – original draft, Writing – review & editing.

## Acknowledgements

We thank the staff of JIRCAS, M. Toyoshima, Y. Nakamura, J. Baba, Y. Fukui, I. Gejima, Y. Takiguchi, K. Ozawa, Y. Masamura, Y. Shirai, Y. Saito, N. Hisatomi, Y. Nonoue, A. Aoyama, T. Nada, M. Karasawa, N. Ohmiya, W. Kawakami, and Y. Kida for their excellent technical assistance. We also thank Y. Nagatoshi, M. Bonifacio, M. Fujita, J. Tanaka, and T. Egi for their support and valuable discussions.

## Generative AI Statement

Generative artificial intelligence tools were used only to assist with language editing and improvement of clarity. The authors reviewed and edited all AI-assisted text and take full responsibility for the content of the submitted manuscript.

## Supplementary material

**Supplementary Table 1**

Taxonomic classification and cube-based allocation of the 135 bacterial isolates used for SynCom construction.

**Supplementary Table 2**

Bioassay-based screening of 45 quinoa-derived SynComs under 100 mM NaCl.

**Supplementary Table 3**

Taxonomic characteristics of the nine bacterial isolates constituting SCDY1.

**Supplementary Table 4**

Primers used for RT-qPCR analysis.

**Supplementary Table 5**

Plant growth-promoting traits of SCDY1 and its constituent bacterial isolates.

**Supplementary Table 6**

Re-isolation of SCDY1-derived bacteria from inoculated *Arabidopsis* seedlings under salinity stress.

**Supplementary Table 7**

Summary of upregulated and downregulated DEGs in the four pairwise transcriptome comparisons.

**Supplementary Table 8**

SCDY1-specific DEGs under salinity stress.

**Supplementary Table 9**

Gene Ontology enrichment analysis of SCDY1-specific DEGs under salinity stress.

**Supplementary Table 10**

Hierarchical clustering of DEG expression patterns across the four treatment groups.

**Supplementary Table 11**

Gene Ontology enrichment analysis of SCDY1-responsive DEGs: biological process.

**Supplementary Table 12**

Gene Ontology enrichment analysis of SCDY1-responsive DEGs: molecular function.

**Supplementary Table 13**

KEGG pathway enrichment analysis of SCDY1-responsive DEGs under salinity stress.

**Supplementary Table 14**

SCDY1-responsive DEGs assigned to the phenylpropanoid biosynthesis pathway.

**Supplementary Table 15**

SCDY1-responsive DEGs assigned to the glutathione metabolism pathway.

